# A pain-reducing Kv6.4 variant spares other Kv6 channels, offering a target for uterine pain

**DOI:** 10.64898/2025.12.18.692915

**Authors:** Ichrak Drissi, Wen Hui Ng, Samiha S Shaikh, Aidan Nickerson, Emanuele Sher, Achim Kless, C Geoffrey Woods

## Abstract

Uterine pain conditions such as dysmenorrhea and endometriosis are highly prevalent, poorly managed, and associated with long-term impacts on women’s health. The identification of a rare *KCNG4* variant (rs140124801; p.Val419Met) previously linked to reduced labor pain suggests Kv6.4 (encoded by *KCNG4*) may play a role in visceral nociception and offer a new target for non-opioid uterine pain relief.

We analyzed UK Biobank data to evaluate clinical phenotypes associated with rare single nucleotide polymorphisms (SNPs) in the conserved TVGYG selectivity filter motif of the Kv6 family wherein p.Val419Met is located. Functional consequences of these variants were assessed using immunofluorescence in SHSY5Y cells to examine membrane trafficking, co-immunoprecipitation to investigate interactions between Kv6 subunits and Kv2.1, and single-cell RNA sequencing to determine expression patterns in mouse sensory neurons.

Our genetic analysis identified 5,816 individuals heterozygous and 28 homozygous for the p.Val419Met variant, as well as 292 heterozygous carriers of the p.Thr418Met variant, all located in the Kv6.4 subunit. Neither variant was associated with increased risk for general, neurological, or pain-related disorders, even in homozygous p.Val419Met carriers, supporting a favorable safety profile. Kv6.4Val419Met has a dominant negative effect on wild type Kv6.4. However, this effect is specific to Kv6.4 as, in SHSY5Y cells, co-expression of Kv6.4Val419Met along with Kv6.1, Kv6.2 or Kv6.3, showed no effects on the efficient membrane localization of Kv6.1-3. In contrast, Kv6.4Thr418Met does not interfere with Kv6.4 trafficking or its heteromerization with Kv2.1. Additionally, In SHSY5Y cells, the equivalent p.Val419Met substitution, when introduced into Kv6.1, Kv6.2 and Kv6.3, disrupts their membrane localization, noting that these variants have never been reported. Co-immunoprecipitation shows that Kv6.4 does not interact with other Kv6 subunits and transcriptomic analysis shows that Kv6.4 is expressed in a distinct subset of mouse lumbar dorsal root ganglion neurons innervating pelvic organs.

Our findings show that the Kv6.4Val419Met variant selectively disrupts Kv6.4 function without affecting other family members and is not linked to adverse phenotypes. This finding supports Kv6.4 as a highly selective and functionally distinct Kv6 subunit with no widespread deleterious effects on other Kv6 subunits, making it an attractive candidate for therapeutic targeting for uterine pain.

## Introduction

Female pain remains historically under-researched and frequently under-treated. Many chronic primary pain conditions, including migraine, and fibromyalgia, exhibit striking sex differences, with women significantly more frequently affected than men.^1^ When including uterine-specific pain conditions such as dysmenorrhea and endometriosis, the cumulative burden significantly reduces women’s quality of life and imposes substantial socioeconomic costs.^2,3^

Dysmenorrhea, or menstrual pain, is one of the most common forms of visceral pain, affecting up to 90% of women of reproductive age, with 10–20% experiencing pain that significantly disrupts daily life.^3,4,5^ Endometriosis, a chronic inflammatory condition affecting approximately 10% of women globally, also can cause debilitating pelvic pain, as well as dysmenorrhea, and infertility.^6^ Labor pain, although acute, is among the most intense forms of human pain. It is typically managed with epidural analgesia, which, while effective, carries potential risks including maternal hypotension, motor blockade, and prolonged labor duration.^7^

In 2020, a rare genetic variant was identified associated with reduced labor pain sensitivity the single nucleotide polymorphism (SNP) rs140124801 in the *KCNG4* gene.^8^ This was reported in a cohort of healthy women who did not request analgesia during their first delivery, despite having normal sensory thresholds.

*KCNG4*, part of the *KCNG* family (*KCNG1–KCNG4*), encodes the Kv6.4 subunit, a silent voltage-gated potassium channel subunit. It modulates neuronal excitability through heteromerization with Kv2 subunits, particularly Kv2.1 (encoded by *KCNB1*), a key delayed rectifier potassium channel expressed in both the central and peripheral nervous systems.^9,10^ While Kv2.1 can form functional homotetrameric channels, it also assembles with silent subunits such as Kv6.4 in a fixed stoichiometry of two Kv2.1 subunits and two Kv6.4 subunits, giving rise to heterotetrameric channels with distinct biophysical properties, compared to the homotetramer.^11,12^

The rs140124801 variant (Kv6.4Met419) disrupts Kv6.4 trafficking to the plasma membrane by impairing the interaction between Kv6.4 and Kv2.1. This prevents Kv6.4 from modulating Kv2.1 function, producing a significantly more depolarized voltage dependence of inactivation of Kv2.1 and ultimately reducing sensory neuron excitability and reducing labor pain.^8^

In this study, we examine whether *KCNG4* represents a safe and selective therapeutic target for uterine and cervix pain, with particular focus on whether Kv6.4Met419 variant impacts the trafficking of other Kv6 family members.

## Materials and methods

### Genetic analysis from the UK biobank data

We analyzed phenotypes associated with *KCNG1-4* single nucleotide variants (SNVs). The analysis focused on the most common disorders in the overall UKBB population (N=446814), as well as in individuals carrying rs140124801 and rs140682724 in the *KCNG4* gene. Particular attention was given to pain-related and neurological phenotypes. The UK Biobank cohort presents a limitation for studying uterine pain in women of reproductive age as it only includes individuals aged 54 to 84.

### DNA constructs

The following wild-type constructs were used in this study: Kv6.1-HA, Kv6.2-Myc, Kv6.3-His, and Kv2.1-IRES-EGFP. Full-length coding sequences of *KCNG1*, *KCNG2*, and *KCNG3* were cloned into pcDNA3.1 vectors with HA, Myc, and His tags, respectively, in the C-terminus. *KCNG1*, *KCNG2*, and *KCNG3* were cloned also into pcDNA3.1 vectors with a Flag tag in the C-terminus. *KCNB1* was cloned into pCAGGS-IRES2-nucEGFP vector. All vectors conferred ampicillin resistance. Constructs were purchased from GenScript.

### Site directed mutagenesis

rs140124801 mutations were introduced into the wildtype *KCNG1-4* plasmids and rs140682724 in *KCNG4* using the QuikChange II XL Site- Directed Mutagenesis Kit (Agilent Technologies). The online QuikChange Primer Design Program was used to generate suitable mutagenesis primers. Mutagenesis and Sanger sequencing primers are listed in Supplementary Fig S1.

### Cell lines and culture conditions

SHSY5Y cells were cultured in 90% Dulbecco’s modified Eagle medium (DMEM) supplemented with 10% fetal bovine serum (FBS), 100 U/ml penicillin-streptomycin, and 2 mM L-glutamine at 37°C, 5% CO_2_, 100% humidity. Transfections were carried out using FugeneHD transfection reagent (Promega) according to the manufacturer’s protocols. For co-expression studies, cloned Kv6.4 with either Kv6.1, Kv6.2, Kv6.3, Kv6.4 or Kv2.1 constructs were co-transfected at a ratio of 1:1except for Kv2.1 where the transfection ratio is 1:2.

### Immunofluorescence analysis and confocal microscopy

1 10^5^ SHSY-5Y cells were cultured on poly-L-lysine coated coverslips and transfected as described above. Un-transfected cells were used as negative controls to assess background autofluorescence. 48-hours after transfection, cells were fixed by 15minutes emersion in ice cold methanol. Cells were permeabilized by 10 minutes incubation in 0.3% Triton X-100 solution followed by 30 minutes blocking in a 10% goat serum blocking buffer (Invitrogen) at room temperature. cells were then stained with primary antibodies overnight at 4°C: Mouse anti-HA (Biolegend, 1:1000), rat anti-Flag (Biolegend, 1:1000) and rabbit anti NaLJ/KLJ-ATPase (Abcam, 4:1000). Secondary antibody used are Alexa Fluor 488 goat anti rat, Alexa Fluor 546 goat anti-mouse, and Alexa Fluor 633 goat anti rabbit (all from Life Technology, 1:1000). Coverslips were mounted onto glass slides using Prolong Diamond Antifade Mountant with DAPI (Molecular Probes). Cells were visualized with an LSM880 confocal microscope. Colocalization between Kv6 subunits and NaLJ/KLJ-ATPase was assessed using the Pearson correlation coefficient as previously described.^8^

### Co-immunoprecipitation and western blot

1.5 10^6^ SHSY-5Y cells were plated onto a 100mm dish and transfected with Kv6.4-HA along with Kv6.1-Flag, Kv6.2-Flag, Kv6.3-Flag or Kv2.1 and harvested after 2 days. Co-immunoprecipitation was carried out using the Dynabeads Co-Immuniprecipitation Kit (Life Technologies) according to the manufacturer’s protocols. 5 µg of an anti-HA antibody (Biolegend) was used per 1 mg of Dynabeads® M-270 Epoxy. Immunoprecipitation buffer supplied in the kit was supplemented with 80mM NaCl. The antibody-coupled DynabeadsR Epoxy was incubated with the cell lysate overnight at 4 °C. Eluted samples were mixed with SDS sample buffer supplemented with dithiothreitol (DTT) and heated at 95 °C for five minutes then loaded onto Bolt™ 4-12% Bis-Tris Plus Gels (Invitrogen, Carlsbad, CA, USA) and separated by SDS-PAGE using MES SDS Running Buffer (Invitrogen). Proteins were subsequently transferred to PVDF membranes using the iBlot™ 2 Dry Blotting System (Invitrogen). Membranes were blocked in 5% (w/v) non-fat dry milk prepared in PBS containing 0.1% Tween-20 (PBST) for one hour at room temperature (RT). Following blocking, membranes were incubated overnight at 4°C with primary antibodies: anti-Flag (BioLegend, 1:1000) and anti-Kv2.1 rabbit polyclonal (Alomone, 1:500). After three washes with PBST, membranes were incubated for 45 minutes at RT with appropriate HRP-conjugated secondary antibodies. Blots were then washed three more times with PBST, developed using enhanced chemiluminescence (ECL) substrate (Cytiva Amersham), and imaged using a ChemiDoc™ Imaging System (Bio-Rad Laboratories).

### Statistical analysis

Statistical analyses were performed using GraphPad Prism version 10 (GraphPad Software, San Diego, CA, USA). Statistical significance from the UKBB analysis was determined using the Monte Carlo version of Fisher’s exact test due to small sample sizes. Statistical significance for Pearson colocalization analysis was determined using a Student’s t-test. The null hypothesis was rejected at a significance level of *p = 0.05*.

## Results

### Variants in the *KCNG* genes from the UKBB population

The Kv6 family contains a conserved ion selectivity filter, the TVGYG motif, wherein the rs140124801 variant is located. This variant results in a p.Val419Met substitution in Kv6.4 (Figure 1A), which has been associated with reduced labor pain.^8^ Based on this finding, we analyzed variants in the TVGYG motif of *KCNG* family subunits across the entire UK Biobank cohort. Notably, *KCNG4* harbors more variants in this motif than other *KCNG* genes (Table1). Specifically, five mutations were identified in *KCNG4*, including *5,816* individuals heterozygous for p.Val419Met, *28* homozygotes for the same variant, and *292* individuals heterozygous for rs140682724 (p.Thr418Met) (Figure1B).

**Table 1.** Variants within the Kv6 subunits’ TVGYG motif identified in the UK Biobank population. SNV: Single nucleotide variant; ID: gnomAD identity; UKBB: UK biobank

**Figure 1.**
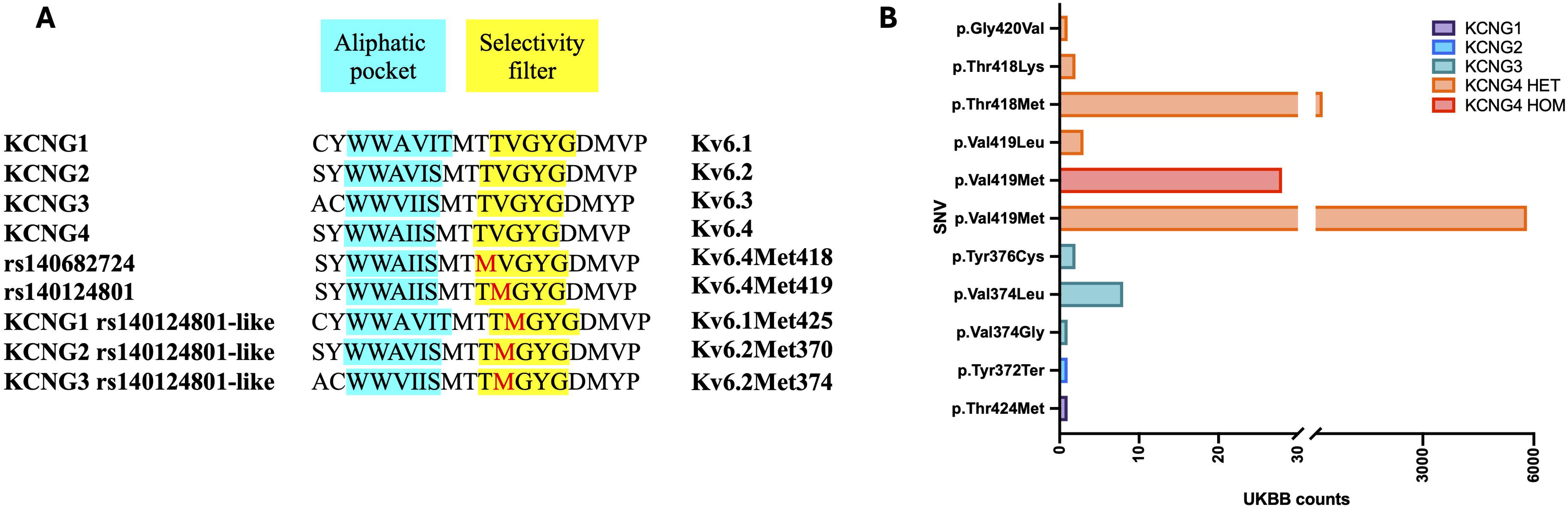
Variant Tolerance in Kv6.4 and the TVGYG Motif. **(A)** Conservation of the selectivity filter TVGYG in human *KCNG1-4* subunits (shown in yellow). rs140682724, rs140124801 and rs140124801-like alleles and representative proteins of each human K_V_6 family member. Invariant amino acids are colored in red. **(B)** Single nucleotide variants (SNVs) identified in the UK Biobank within the TVGYG motif across all *KCNG* genes within a cohort of middle aged (57-87 years old) women and men. *KCNG4* shows the highest number of SNVs in this conserved region compared to other family members, suggesting a higher tolerance for mutation

### Clinical phenotype analysis in the UKBB database

Clinical phenotypes were analyzed in individuals carrying either the heterozygous or homozygous *KCNG4* variant rs140124801, as well as those heterozygous for rs140682724. The analysis included *(i)* common disorders (e.g., hypertension (I20), hypercholesterolemia (E78.0), obesity (E66.9), cataract (H26.9)), *(ii)* pain-related conditions (e.g., chest pain (R07.4), arthrosis (M19.9), headache (R51), sciatica (M54.3)), and *(iii)* neurological disorders (e.g., epilepsy (G40.9), dementia (F03), Alzheimer’s disease (G30.9)). See supplementary material for all the examined phenotypes and their respective codes in the International Classification of Diseases 10^th^ revision (ICD-10) https://icd.who.int/browse10/2019/en (Supplementary Fig S2). We included both women and men from the UK Biobank cohort. First, the frequencies of common disorders, pain-related conditions, and neurological disorders in both heterozygous and homozygous carriers of rs140124801 were compared with those in the general UK Biobank population. No significant differences were found (Monte Carlo Fisher’s exact test, *p > 0.05*), indicating that rs140124801 does not affect the prevalence of these clinical phenotypes (Supplementary Fig. S3). Then, a comparison of clinical phenotype frequencies, including common disorders, pain-related conditions, and neurological disorders, was performed between heterozygous carriers of the rs140124801 and rs140682724 variants and the general UK Biobank population. No significant differences were observed among the three groups (Monte Carlo Fisher’s exact test, *p>0.05*) (Supplementary Fig S4).

Although women in the UK Biobank cohort are generally beyond reproductive age, we specifically analysed uterine-related conditions (endometriosis, dyspareunia, and excessive menstruation) in female heterozygous carriers of the rs140124801 and rs140682724 variants, compared to the broader female UKBB population. No significant differences were observed between the groups (Monte Carlo Fisher’s exact test, p > 0.05) (Supplementary Fig S5).

### Membrane localization of Kv6.1, Kv6.2, and Kv6.3 is impaired by rs140124801-like substitutions

As previously discussed, the SNP rs140124801 in the *KCNG4* gene disrupts the interaction between Kv6.4 and Kv2.1, preventing Kv6.4 from reaching the plasma membrane.^8^ This mutation lies within the highly conserved TVGYG motif of the selectivity filter, a region that is preserved across all *KCNG* family members. To assess whether rs140124801-like variants exert similar effects across other *KCNG* family members, we introduced equivalent valine-to-methionine substitutions at position 425 in Kv6.1, position 370 in Kv6.2, and position 374 in Kv6.3, each corresponding to the position 419 in Kv6.4 (Fig1A). Subcellular localization of the resulting variants was then examined by immunofluorescence. Silent potassium subunits are known to be retained in the endoplasmic reticulum in the absence of Kv2.1.^9,10^ Therefore, we used SH-SY5Y cells, which endogenously express Kv2.1 shown by real time PCR (Supplementary methods and Supplementary Fig S6).

SHSY5Y cells expressing Kv6.1-HA wild-type (WT) showed efficient trafficking of Kv6.1 to the plasma membrane, however the Kv6.1Met425-HA mutant failed to reach the plasma membrane and was retained intracellularly (Figure 2A). Pearson colocalization analysis between Kv6.1-HA and NaLJ/KLJ-ATPase revealed a significant reduction in the colocalization coefficient for the Kv6.1Met425 variant and NaLJ/KLJ-ATPase compared to Kv6.1 WT (*p<0.0001*), indicating impaired membrane localization (Figure 2D).

**Figure 2.**
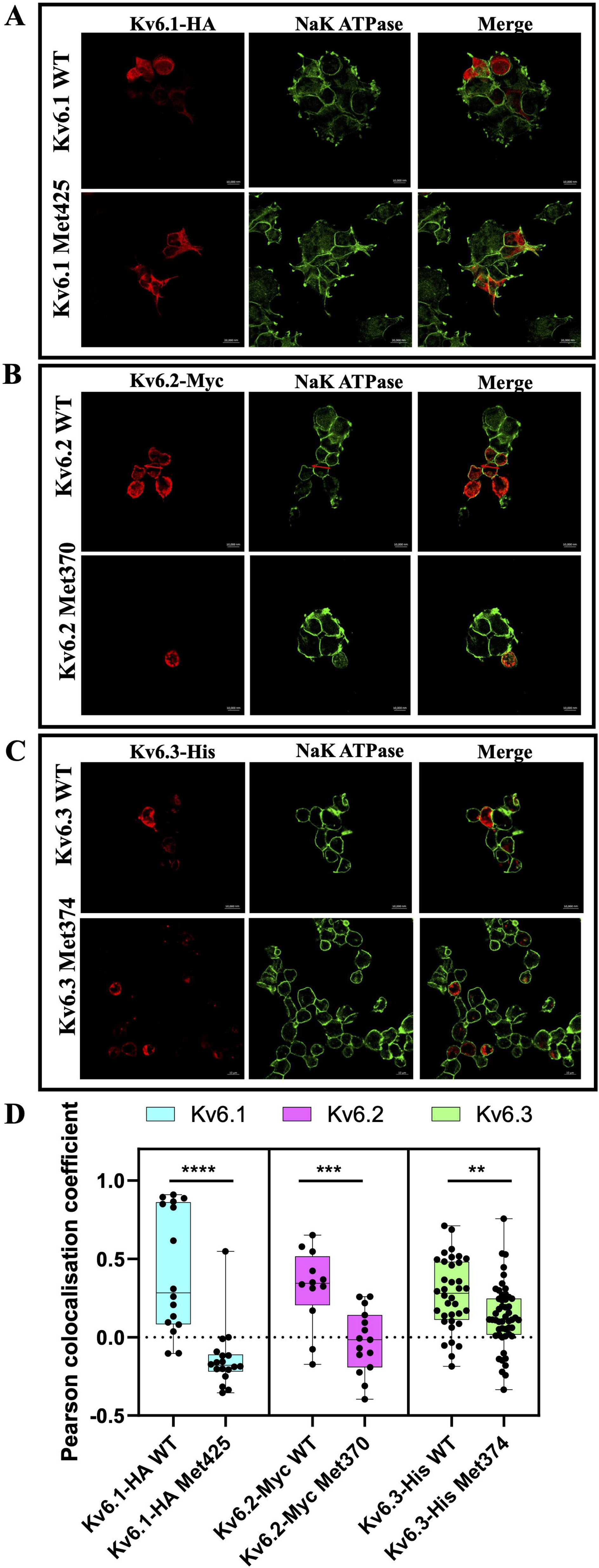
rs140124801-like variants impair membrane localization of Kv6.1, Kv6.2, and Kv6.3 subunits. SH-SY5Y cells were transfected with either Kv6.1-HA wild-type, Kv6.1Met425-HA, Kv6.2-Myc wild-type, Kv6.2Met370-Myc, Kv6.3-His wild-type, or Kv6.3Met374-His constructs. Transfected cells were co-stained using anti-HA, anti-Myc, or anti-His antibodies (red), with NaLJ/KLJ-ATPase (green) as a plasma membrane marker. **(A)** Co-immunostaining of Kv6.1-HA wild-type (top panel) and Kv6.1Met425-HA (second panel), co-stained with the NaLJ/KLJ-ATPase. **(B)** Co-immunostaining of Kv6.2-Myc wild-type (top panel) and Kv6.2Met370-Myc (second panel) co-stained with the NaLJ/KLJ-ATPase. **(C)** Co-immunostaining of Kv6.3-His wild-type (top panel) and Kv6.3Met374-His (second panel) co-stained with the NaLJ/KLJ-ATPase. **(D)** Pearson colocalization coefficients between each Kv6 subunit and NaLJ/KLJ-ATPase were quantified to assess membrane localization. The box-and-whisker plot displays Pearson correlation values for each condition, with each dot representing an individual stained cell. Experiments were independently repeated four times (*N = 4*). Statistical differences between wild-type and corresponding mutant constructs were evaluated using Student’s *t*-test (*\*\*p < 0.01, ***p < 0.001*).

A similar pattern was observed for the Kv6.2Met370 variant. Immunofluorescence imaging of SH-SY5Y cells transfected with either Kv6.2-Myc WT or Kv6.2Met370-Myc, followed by Pearson colocalization analysis with NaLJ/KLJ-ATPase, showed a significant decrease in membrane colocalization for the Kv6.2Met370 variant compared to wild-type Kv6.2 (*p<0.001*) (Figure 2B, 2D).

Likewise, the Kv6.3Met374 variant showed a reduced plasma membrane localization relative to Kv6.3 WT, as demonstrated by both immunofluorescences staining and a significant decrease in Pearson’s correlation coefficient with NaLJ/KLJ-ATPase (*p<0.01*) (Figure 2C, 2D).

### The dominant-negative effect of the Kv6.4Met419 variant is subunit-specific, affecting only Kv6.4

To investigate whether the dominant-negative effect of Kv6.4Met419, previously reported,^8^ also impacts other Kv6 family members. Kv6.4-HA WT or Kv6.4Met419-HA, with Kv6.1-Flag WT, Kv6.2-Flag WT, or Kv6.3-Flag WT were expressed in SHSY5Y cells and immunostained with anti-HA, anti-Flag, and NaLJ/KLJ-ATPase antibodies.

Figure 3A shows immunostaining of SH-SY5Y cells co-transfected with Kv6.4-HA WT or Kv6.4Met419-HA, together with Kv6.1-Flag WT. Pearson colocalization analysis between Kv6.1-Flag WT and NaLJ/KLJ-ATPase revealed a reduction in the colocalization coefficient in the presence of the Kv6.4Met419 variant compared to Kv6.4 WT (*p<0.05*). However, the Pearson coefficient remained positive, indicating that Kv6.1 retains its ability to reach the plasma membrane even when co-expressed with the rare Kv6.4Met419 variant (Figure 3A, 3D).

**Figure 3.**
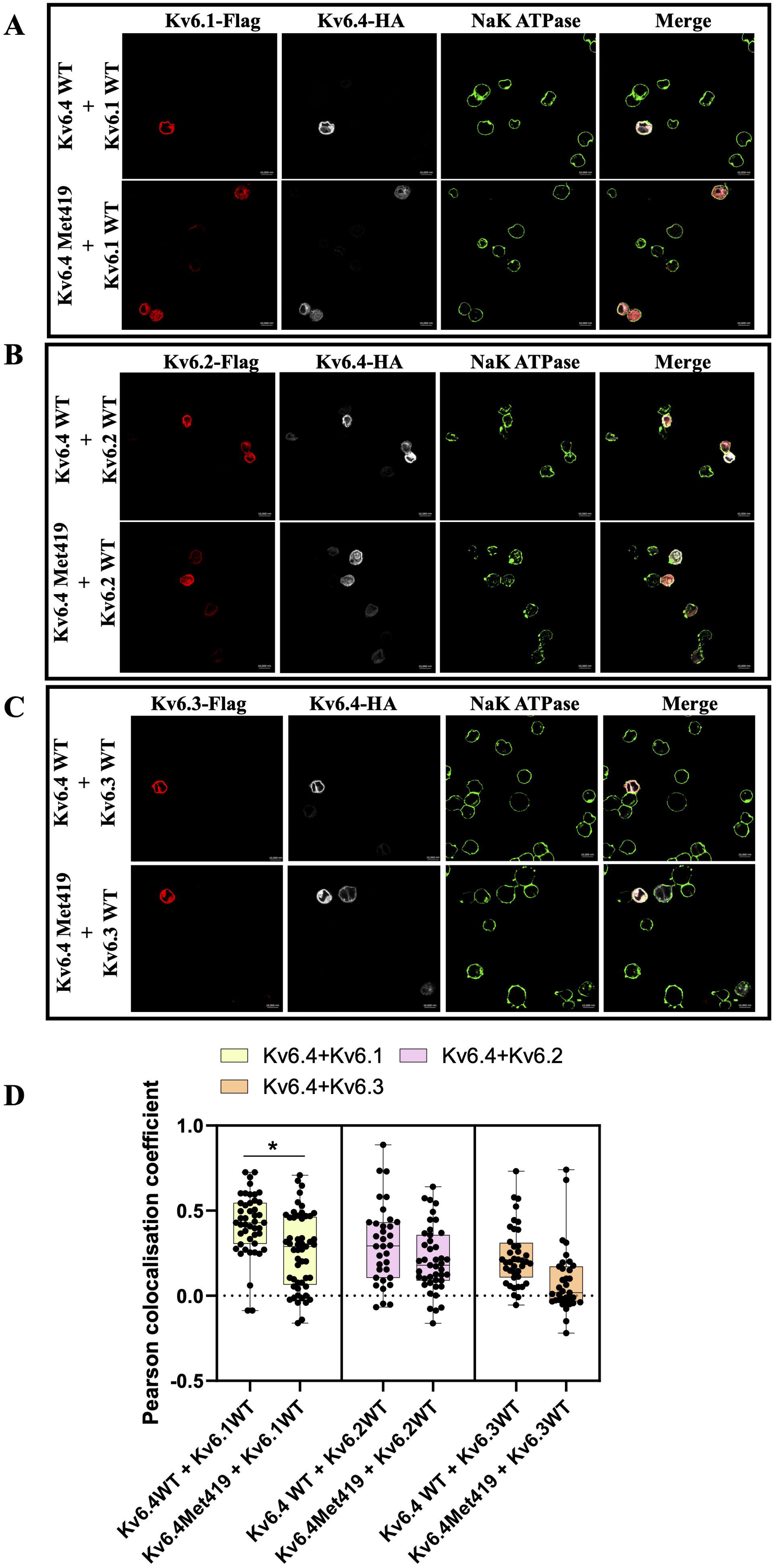
The dominant-negative effect of Kv6.4Met419 is specific to Kv6.4 subunit. SH-SY5Y cells were co-transfected with either wild-type Kv6.4-HA or the Kv6.4Met419-HA mutant (both shown in white), along with wild-type Kv6.1-Flag, Kv6.2-Flag, or Kv6.3-Flag (shown in red). Cells were co-stained with anti-HA and anti-Flag antibodies, as well as NaLJ/KLJ-ATPase (green) to mark the plasma membrane. **(A)** Co-immunostaining of Kv6.4-HA wild-type (top panel) or Kv6.4Met419-HA (second panel) with Kv6.1-Flag and NaLJ/KLJ-ATPase. **(B)** Co-immunostaining of Kv6.4-HA wild-type (top panel) or Kv6.4Met419-HA (second panel) with Kv6.2-Flag and NaLJ/KLJ-ATPase. **(C)** Co-immunostaining of Kv6.4-HA wild-type (top panel) or Kv6.4Met419-HA (second panel) with Kv6.3-Flag and NaLJ/KLJ-ATPase. **(D)** Pearson colocalization coefficients between Flag-tagged subunits and NaLJ/KLJ-ATPase were quantified to assess membrane localization. Kv6.1, Kv6.2, and Kv6.3 retained effective membrane localization when co-expressed with Kv6.4Met419. The box-and-whisker plot displays Pearson correlation values for each condition, with each dot representing an individual stained cell. Experiments were independently repeated four times (*N = 4*). Statistical comparisons between wild-type and mutant co-expression conditions were performed using Student’s t-test (*\*p < 0.05*).

Co-expression of Kv6.2-Flag WT with either Kv6.4-HA WT or Kv6.4Met419-HA (Figure 3B) followed by immunostaining and Pearson colocalization analysis showed no significant difference in membrane localization of Kv6.2, confirming that it efficiently traffics to the plasma membrane in both conditions (*p>0.05*) (Figure 3B, 3D)

Likewise, co-expression of Kv6.3-Flag WT with either Kv6.4-HA WT or Kv6.4Met419-HA in SH-SY5Y cells showed no alteration in Kv6.3 membrane localization. Immunostaining and Pearson colocalization analysis revealed comparable coefficients between Kv6.3-Flag and NaLJ/KLJ-ATPase in the presence of the Kv6.4Met419 variant (*p>0.05*), indicating that Kv6.3 membrane trafficking is unaffected by Kv6.4Met419 (Figure 3C, 3D).

### Kv6.4 membrane localization and interaction with Kv2.1 remain unaffected by the rs140682724 variant

As described above, analysis of the UK Biobank database identified 292 individuals carrying the rs140682724 variant. This SNP results in an amino acid substitution at position 418 in Kv6.4, changing threonine to methionine. Since Thr418 lies within the conserved TVGYG selectivity filter motif, we investigated whether rs140682724 (Kv6.4Thr418Met) affects Kv6.4 subcellular localization and its interaction with Kv2.1. Additionally, we tested whether the Kv6.4Thr418Met variant reduces interaction with Kv2.1, potentially impairing Kv6.4 trafficking, similar to the effect previously observed for Kv6.4-419Met.^8^

SH-SY5Y cells were transfected with either Kv6.4-HA WT or Kv6.4Thr418Met-HA and immunostained with anti-HA and NaLJ/KLJ-ATPase antibodies (Figure 4A). Pearson correlation coefficient showed no significant difference between Kv6.4-HA WT and Kv6.4Met418-HA (*p > 0.05*), with both conditions showing a positive Pearson coefficient, indicating effective membrane localization of Kv6.4Met418 (Figure 4B).

**Figure 4:**
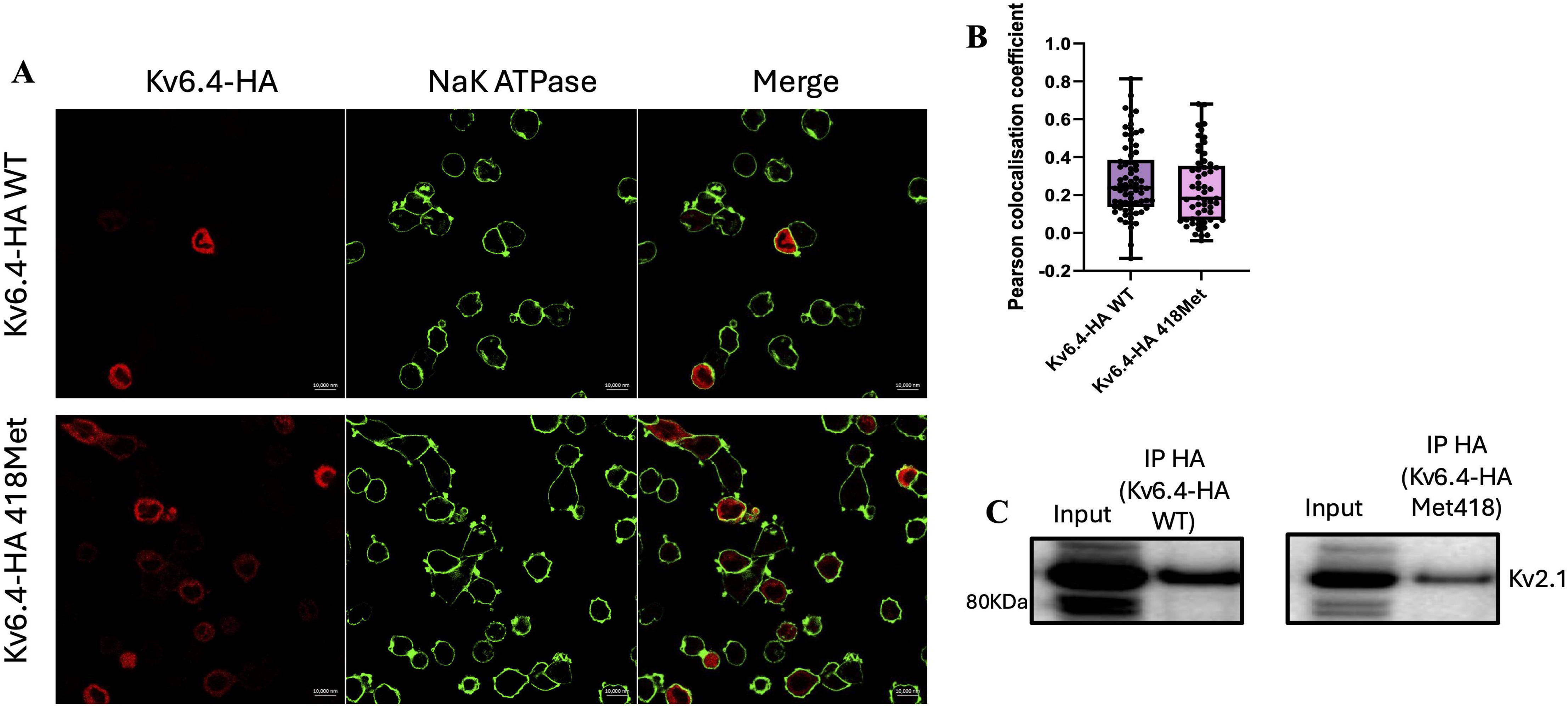
Kv6.4Met418 preserves subcellular localization and interaction with Kv2.1, unlike Kv6.4Met419. SH-SY5Y cells were transfected with either Kv6.4-HA wild-type, Kv6.4-Met418-HA and immnuostained with anti-HA antibody (red) and NaLJ/KLJ-ATPase (green). **(A)** Immunostaining of Kv6.4-HA wild-type (top panel) or Kv6.4Met418-HA (second panel) and NaLJ/KLJ-ATPase. **(B)** Pearson colocalization coefficients between HA-tagged Kv6.4 and NaLJ/KLJ-ATPase were analyzed to assess membrane localization. Kv6.4Met418 analysis shows a Pearson colocalization coefficient with NaLJ/KLJ-ATPase comparable to that of Kv6.4 WT, indicating that Kv6.4Met418 successfully reaches the plasma membrane. The box-and-whisker plot displays Pearson correlation values for each condition, with each dot representing an individual stained cell. Experiments were independently repeated four times (*N = 4*). Statistical comparisons between wild-type and mutant co-expression conditions were performed using Student’s t-test. (**C)** Co-immunoprecipitation of Kv6.4-HA or Kv6.4-HAMet418 using an anti-HA antibody. Following pulldown, Kv2.1 (95 kDa) was detected using an anti-Kv2.1 antibody. The results show that Kv6.4Met418 interacts with Kv2.1 similarly to wild-type Kv6.4, although interaction strength varied between experiments. The blot has been cropped; full-length blots are provided in Supplementary Fig S7D. *Input = lysate from transfected cells; IP = co-immunoprecipitated sample.

SH-SY5Y cells were transfected with either Kv6.4-HA WT or Kv6.4Thr418Met-HA, and Kv6.4 was immunoprecipitated using an anti-HA antibody. Co-immunoprecipitation assays demonstrated that Kv6.4Met418 retains the ability to interact with Kv2.1 (Figure 4C).

### Kv6.4 is expressed in a distinct sensory neuron subset and does not interact with other Kv6 subunits

To assess whether Kv6.4 could serve as a specific therapeutic target for female-specific pain without interfering with other Kv6 subunits, we examined its potential interactions with Kv6 subunits. Kv6.4-HA WT was co-expressed with Kv6.1-Flag WT, Kv6.2-Flag WT, Kv6.3-Flag WT, or Kv2.1 (positive control), and immunoprecipitated using an anti-HA antibody.

Co-immunoprecipitation revealed that Kv6.4 does not interact with any other Kv6 subunits but does interact with Kv2.1, as expected (Figure 5A).

**Figure 5:**
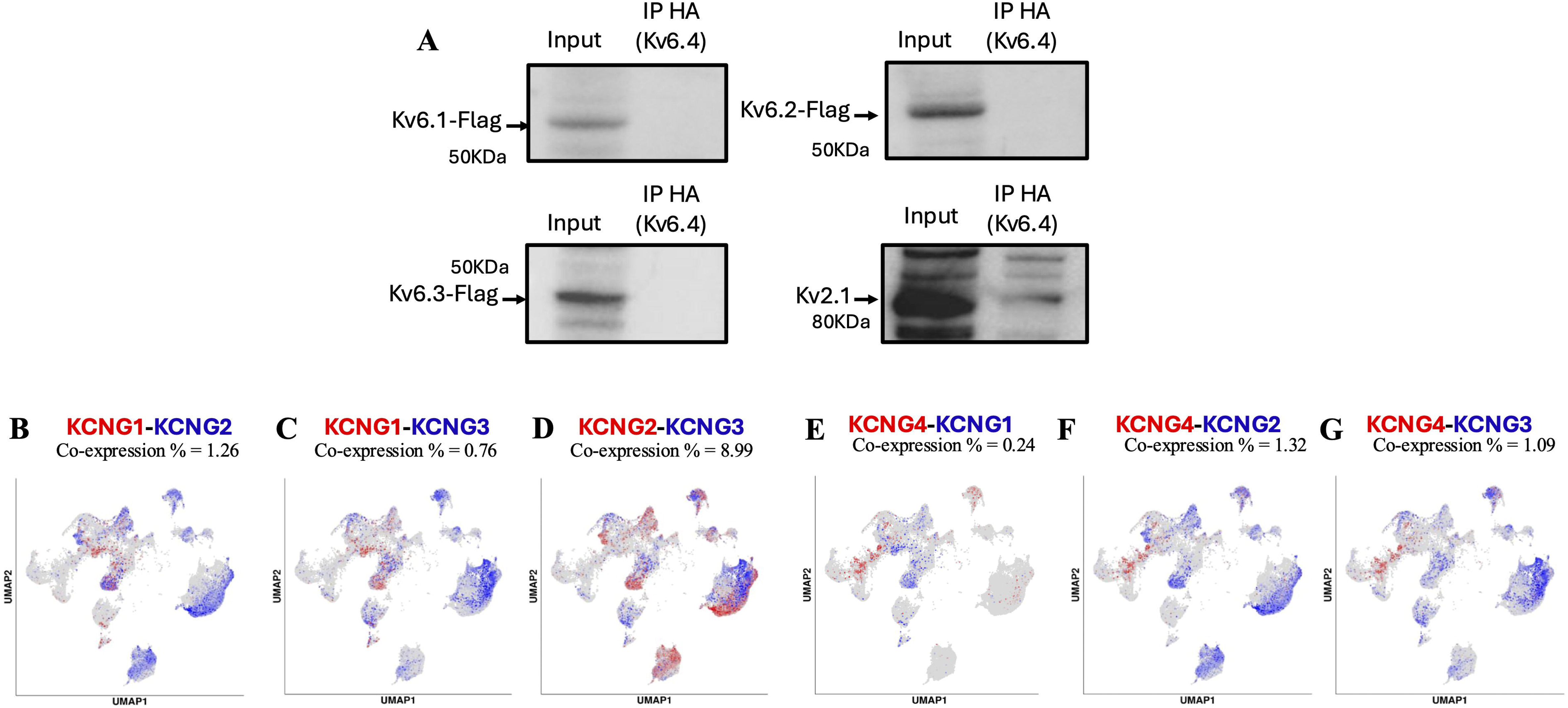
Kv6.4 neither interacts with nor colocalizes with other Kv6 subunits. **(A) Kv6.4 does not interact with other Kv6 subunits:** SH-SY5Y cells were co-transfected with Kv6.4-HA (wild-type) and either Kv6.1-Flag, Kv6.2-Flag, Kv6.3-Flag, or Kv2.1 (used as a positive control). Following immunoprecipitation of Kv6.4-HA using an anti-HA antibody, co-immunoprecipitated proteins were probed using an anti- Flag antibody for Kv6.1, Kv6.2, and Kv6.3, and a Kv2.1-specific antibody. The co-immunoprecipitation results shows that Kv6.4 does not interact with Kv6.1, Kv6.2, or Kv6.3, but does interact with Kv2.1 as expected. Cropped blot shown; the full-length blot is available in Supplementary Fig S7. *Input = lysate from transfected cells; IP = co-immunoprecipitated sample. **(B) Expression pattern of *KCNG4* and other *KCNG* subunits in mouse sensory neurons:** Single-cell analysis reveals that *KCNG4* is expressed in a distinct population of DRG sensory neurons and does not colocalize with other *KCNG* subunits (Analysis based on Bhuiyan et al., 2024) (http://harmonized.painseq.com)

We next examined the expression patterns of all *KCNG* family members in mouse sensory neurons using the PainSeq single cell database https://painseq.shinyapps.io/harmonized_painseq_v1/.^14^ Single-cell analysis revealed low co-expression rates among *KCNG* subunits: *KCNG1–KCNG2* (1.26%), *KCNG1–KCNG3* (0.76%), and *KCNG2–KCNG3* (8.99%, highest among pairs). *KCNG4* showed minimal co-expression with *KCNG1* (0.24%), *KCNG2* (1.32%), and *KCNG3* (1.09%), suggesting that *KCNG4* is expressed in a distinct subset of sensory neurons (Figure 5B). However, these databases are enriched for lumbar DRGs, which is useful for our work as these innervate the uterus, but are poor for thoracic and sacral DRG where thoracolumbar (T10-L2) and sacral (S2-4) DRGs serve the cervix.

### The rs140124801-like variant in *KCNG1* does not disrupt the localization of Kv6.3 and Kv6.2 nor does not exert a dominant-negative effect on WT Kv6.1

As reported above, the rs140124801-like variant disrupts the subcellular localization of Kv6.1, Kv6.3, and Kv6.4, mirroring the effect of rs140124801 on Kv6.4. Since the dominant-negative effect of Kv6.4Met419 appears to be specific to the Kv6.4 subunit, we investigated whether the rs140124801-like variant in Kv6.1 (Kv6.1Met425) could similarly act in a dominant-negative manner or interfere with the localization of other Kv6 subunits.

Figure 6A shows immunostaining of SH-SY5Y cells co-transfected with Kv6.1-HA WT or Kv6.1Met425-HA and Kv6.1-Flag WT to assess potential dominant-negative effects. Pearson colocalization analysis between Kv6.1-FlagWT and NaLJ/KLJ-ATPase revealed no significant difference in membrane localization in the presence of Kv6.1Met425 compared to Kv6.1 WT (*p > 0.05*; Figure 6D). Similarly, co-expression of Kv6.1-HA WT or Kv6.1Met425 with either Kv6.2-Flag WT or Kv6.3-Flag WT (Figures 6B and 6C) also showed no impairment in plasma membrane localization. Pearson colocalization coefficients confirmed that Kv6.2 and Kv6.3 reached the membrane at levels comparable to when co-expressed with Kv6.1 WT (*p > 0.05*; Figure 6D).

**Figure 6:**
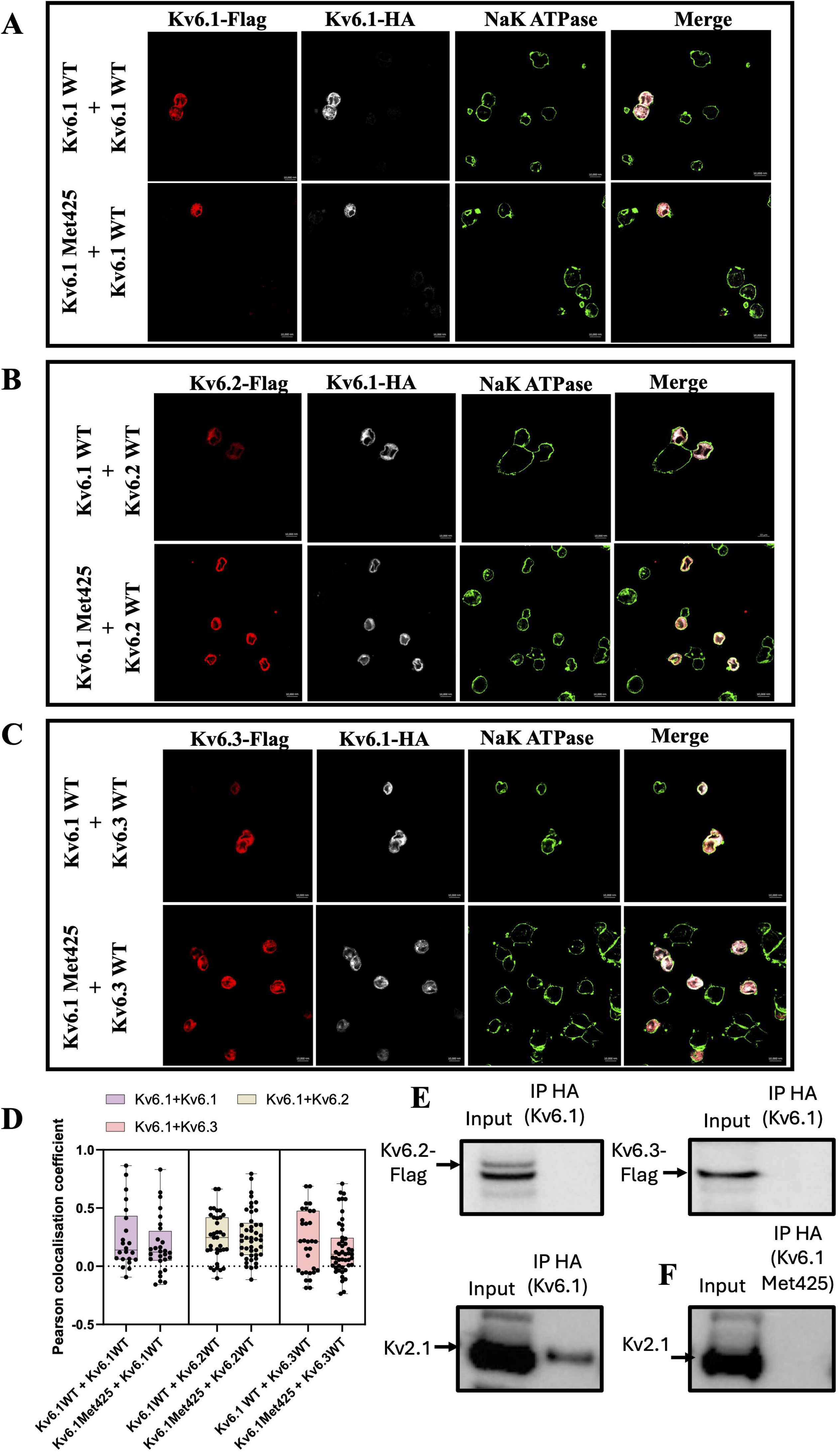
Kv6.1Met425 does not act as a dominant negative. SH-SY5Y cells were co-transfected with either wild-type Kv6.1-HA wild type or the Kv6.4Met425-HA mutant (both shown in white), along with wild-type Kv6.1-Flag wild type, Kv6.2-Flag wild type, or Kv6.3-Flag wild type (shown in red). Cells were co-stained with anti-HA and anti-Flag antibodies, as well as NaLJ/KLJ-ATPase (green) to mark the plasma membrane. (**A)** Co-immunostaining of Kv6.1-HA wild-type (top panel) or Kv6.1Met425-HA (second panel) with Kv6.1-Flag wild type and NaLJ/KLJ-ATPase. **(B)** Co-immunostaining of Kv6.1-HA wild-type (top panel) or Kv6.1Met425-HA (second panel) with Kv6.2-Flag wild type and NaLJ/KLJ-ATPase. **(C)** Co-immunostaining of Kv6.1-HA wild-type (top panel) or Kv6.1Met425-HA (second panel) with Kv6.3-Flag wild type and NaLJ/KLJ-ATPase. **(D)** Pearson colocalization coefficients between Flag-tagged Kv6 subunits and NaLJ/KLJ-ATPase were quantified to assess membrane localization. Kv6.1, Kv6.2, and Kv6.3 maintained effective membrane localization when co-expressed with Kv6.1Met425. The box-and-whisker plot displays Pearson correlation values for each condition, with each dot representing an individual stained cell. Experiments were independently repeated four times (*N = 4*). Statistical comparisons between wild-type and mutant co-expression conditions were performed using Student’s t-test. **(E)** SH-SY5Y cells were co-transfected with Kv6.1-HA (WT) and either Kv6.2-Flag (WT) or Kv6.3-Flag (WT). Kv6.1 was immunoprecipitated using an anti-HA antibody, and the presence of co-immunoprecipitated Kv6.2 or Kv6.3 was assessed using an anti-Flag antibody. Co-immunoprecipitation results show that Kv6.1 does not interact with either Kv6.2 or Kv6.3. A weak interaction between Kv6.1 and Kv6.2 was observed in a single replicate (data not shown). **(F)** SH-SY5Y cells expressing Kv6.1Met425-HA were subjected to co-immunoprecipitation using an anti-HA antibody, and the presence of Kv2.1 was assessed using an anti-Kv2.1 antibody. The results show that the Met425 variant disrupts the interaction between Kv6.1 and Kv2.1. Cropped blots shown; the full-length blots are available in Supplementary Fig S8 *Input = lysate from transfected cells; IP = co-immunoprecipitated sample.

We next examined whether Kv6.1 interacts with other Kv6 subunits and whether the Kv6.1Met425 variant disrupts heteromerization with Kv2.1. Co-immunoprecipitation showed no interaction between Kv6.1 and either Kv6.2 or Kv6.3. A weak interaction between Kv6.1 and Kv6.2 was observed in a single replicate (data not shown). As a positive control, a strong interaction between Kv6.1 and Kv2.1 was observed (Figure 6E). In contrast, the Kv6.1 Met425 variant failed to heteromerize with Kv2.1, mirroring the previously reported effect of Kv6.4Met419 on Kv6.4–Kv2.1 interaction.^8^ (Figure 6F).

## Discussion

In this study, we provide genetic and functional evidence supporting *KCNG4* as a selective and safe therapeutic target for uterine pain. Building on prior associations between the rare variant rs140124801 (p.Val419Met) and reduced labor pain, our analysis of UK Biobank data revealed no association between Kv6.4Met419 or Kv6.4Met418 and clinical phenotypes related to general diseases, pain, or neurological disorders. Functional assays demonstrated that rs140124801/Kv6.4Val419Met-like variants impair membrane trafficking of Kv6.1, Kv6.2, and Kv6.3, mirroring the trafficking defect observed with Kv6.4Met419; and interestingly, none of these other variants have been reported in population databases. Importantly, the dominant-negative effect of the p.Val419Met mutation was specific to Kv6.4 and did not affect to other Kv6 subunits. Furthermore, the Kv6.4Thr418Met variant did not disrupt trafficking or heteromerization with Kv2.1; suggesting that the Kv6.4Thr418Met variant has unexpectedly unique function. We also show that Kv6.4 neither co-localizes nor interacts with other Kv6 subunits and is expressed in a distinct subset of mouse dorsal root ganglion sensory neurons, further supporting its selectivity as a therapeutic target for uterine and pelvic pain.

Uterine pain, such as dysmenorrhea and endometriosis, remains a pervasive and poorly treated condition with variable response to treatments.^15^ Recently, it has been shown that dysmenorrhea in adolescence adversely affects immediate wellbeing and contributes to an increased risk of chronic pain in adulthood, thus lending supporting evidence to calls to consider adolescent dysmenorrhea a crucial public health issue.^3^ Current pharmacological options, such as nonsteroidal anti-inflammatory drugs (NSAID), are often insufficient and associated with systemic side effects such as gastrointestinal and cardiovascular risks,^16^ underlying the need for more targeted and safer therapeutics. In this study, we evaluated the therapeutic potential and safety profile of targeting *KCNG4* for uterine, and possibly also pelvic, pain relief.

### Is there genetic and functional validation of Kv6.4 as a therapeutic target?

Using data from the UK Biobank, we confirmed the presence of several rare missense variants within the highly conserved TVGYG ion selectivity filter motif of *KCNG* family members. Notably, Kv6.4 shows the highest variant number in this region, suggesting a greater tolerance for mutation. Clinical phenotyping revealed no associations between the Kv6.4Met419 variant and neurological, pain-related, or general medical conditions, even in homozygous carriers. Functional assays demonstrated that Kv6.4 p.Val419Met did not interfere with membrane trafficking of other Kv6 subunits.

Kv6.4 represents a promising new target for uterine and cervix pain. Prior work established that the rs140124801 variant (Kv6.4Met419) is significantly enriched in women experiencing less labor pain. Kv6.4 is expressed in a substantial fraction of uterine-projecting sensory neurons co-expressing Kv2.1 and nociceptor markers such as TRPV1, *SCN10A* (Nav1.8), and *SCN11A* (Nav1.9). The rs140124801 variant impairs Kv6.4 trafficking, resulting in altered voltage dependence of inactivation and elevated action potential thresholds.^8^ In parallel, another *KCNG4* variant (Kv6.4-L360P) associated with migraine also alters Kv6.4/Kv2.1 heteromeric function,^17^ supporting the applicability of the role of Kv6.4 in sensory neuron excitability. Our data extend these findings: we show that other *KCNG4* variants in the conserved TVGYG motif are not linked with adverse clinical phenotypes in the UK Biobank, that only Kv6.4Met419 variant (not Kv6.1Met425) act as a dominant-negative, and that Kv6.4 is expressed in a distinct subset of sensory neurons and does not heteromerize with other Kv6s highlighting its potential as a selective and safe therapeutic target, particularly given that *KCNG2* has been implicated in familial sick sinus syndrome.^18^ Transcriptomic profiling of sacral dorsal root ganglia in adult mice confirmed the expression of *KCNG4* in sensory neurons innervating pelvic organs, including the cervix and uterus.^19^ This regional expression supports Kv6.4’s role in visceral nociception and its potential as a target for female-specific pain modulation. Taken together, these findings argue and support that selective modulation or inhibition of Kv6.4 could provide a safe analgesia for uterine/cervical/pelvic pain.

Analysis of clinical phenotypes related to pelvic and uterine pain in women using the UK Biobank was limited by the demographic composition of the cohort, which predominantly includes middle-aged individuals (ages 57–87). Therefore, women of reproductive age are grossly underrepresented, making it difficult to assess the prevalence of conditions such as endometriosis, dysmenorrhea, and labor pain. These limitations highlight the need to expand the UKBB to include younger, reproductive-age women to enable more comprehensive analysis of female-specific pain phenotypes.

A limitation of this study is that we have not performed patch-clamp recordings, although this has been previously published for Kv6.4Val419Met variant on Kv2.1 channel behavior.^8^ Future studies incorporating direct electrophysiological measurements will be essential to confirm the mechanistic implications on neuronal excitability.

### Therapeutic Translation

While a recent study has demonstrated that combinations of Kv2 inhibitors can distinguish Kv2-only channels from Kv2/Kv silent subunits heteromers, including Kv2/Kv6.4 complexes,^20^ no selective small-molecule modulators of Kv6.4 have yet been identified. This reflects both the silent nature of Kv6 family members and their need for heteromerization with Kv2.1 to form functional channels and exit the endoplasmic reticulum.^9,10,21^ Notably, Kv2 channels, including Kv2.1, also serve a structural role in organizing endoplasmic reticulum–plasma membrane (ER/PM) junctions, and Kv6.4 has been shown to cluster with Kv2.1 at these junctions.²¹LJ²³These observations support the need to explore novel pharmacological strategies specifically targeting Kv6.4/Kv2.1 complexes, as well as a deeper investigation into the mechanisms by which Kv2.1/Kv6.4 heteromers traffic from the endoplasmic reticulum to the plasma membrane. An alternative therapeutic approach could involve silencing Kv6.4 using either a mutant Kv6.4Met419 peptide, which acts as a dominant-negative or transient knockdown via siRNA to prevent uterine and pelvic pain.

## Data availability

The data that support the findings of this study are available from the corresponding author, upon reasonable request.

## Acknowledgements

UK Biobank is a large-scale biomedical database and research resource containing genetic, lifestyle and health information from half a million UK participants. This research has been conducted using data from UK Biobank, a major biomedical database under project ID 64765. The UK Biobank database received ethical approval from the North-West Haydock Research Ethics Committee (REC reference 21/NW/0157) and participants gave informed consent. We would like to thank all participants of the UK Biobank cohort who have provided necessary genetic and phenotypic information.

## Funding

ID, SSS, and CGW acknowledge funding from MRC ADVANTAGE visceral pain consortium (MR/W002426/1). ADVANTAGE is part of the UKRI Advanced Pain Discovery Platform initiative funded by MRC, ESRC, BBSRC, Versus Arthritis, Medical Research Foundation, Eli Lilly, and AstraZeneca. CGW acknowledge funding from the Cambridge BioMedical Research Campus (G122118 Cambridge University Hospitals NHS Foundation Trust).

## Competing interests

A.N. and A.K. are current or former employees of Eli Lilly and Company and may own stock in this company.

## Supplementary material

**Supplementary Fig S1**: Mutagenesis primers and Sanger sequencing primers.

**Supplementary Fig S2**: List of clinical phenotypes examined in this study, along with their corresponding ICD-10 codes.

**Supplementary Fig S3**: Frequencies of clinical phenotypes in female and male carriers of rs140124801 (homozygous (*N=28)* and heterozygous (*N=5816*)) variant compared to the general UK Biobank population (*N=446814*). A: Frequencies of the most common disorders. B: Frequencies of pain-related disorders. C: Frequencies of neurological disorders.

**Supplementary Fig S4**: Frequencies of clinical phenotypes in female and male heterozygous carriers of rs140124801 (*N=5816*), heterozygous carriers of rs140682724 (*N=292*) variants compared to the general UKBB cohort. A: Frequencies of the most common disorders. B: Frequencies of pain-related disorders. C: Frequencies of neurological disorders.

**Supplementary Fig S5:** Frequencies of uterine pain-related clinical phenotypes in female heterozygous carriers of the rs140124801 and rs140682724 variants, compared to the general UK Biobank cohort.

**Supplementary methods:** RT-PCR protocol used to assess *KCNB1* expression levels in SHSY5Y and HEK293 cells.

**Supplementary Fig S6:** *KCNB1* relative expression by RT-PCR in SHSY5Y and HEK293 cells. Statistical differences between HEK293 and SH-SY5Y are indicated as *\*p<0.05* (Student’s T-test).

**Supplementary Fig S7: Full Western blots from co-immunoprecipitation experiments. (A–D)** SH-SY5Y cells were co-transfected with Kv6.4-HA (wild-type) and either Kv6.1-Flag **(A),** Kv6.2-Flag **(B),** or Kv6.3-Flag **(C)** Kv2.1 **(D)**. Following immunoprecipitation of Kv6.4-HA using an anti-HA antibody, the presence of co-immunoprecipitated Kv6 subunits was assessed by immunoblotting with an anti-Flag antibody. Expected molecular weights: Kv6.1 (57 kDa), Kv6.2 (51 kDa), and Kv6.3 (49 kDa). Kv2.1 presence was assessed using an anti Kv2.1 antibody (95kDa). **(D)** SH-SY5Y cells expressing either Kv6.4-HA wild-type or Kv6.4-HAThr418Met were subjected to immunoprecipitation using an anti-HA antibody. Co-immunoprecipitated Kv2.1 was detected by immunoblotting with an anti-Kv2.1 antibody.

*Input = lysate from transfected cells; IP = co-immunoprecipitation product; N= number of replicates

**Supplementary Fig S8: Full Western blots from co-immunoprecipitation experiments.**

**(A)** SH-SY5Y cells expressing either Kv6.1-HA wild-type or Kv6.1-HA Met425 were subjected to immunoprecipitation using an anti-HA antibody. Co-immunoprecipitated Kv2.1 (95 kDa) was detected by immunoblotting with an anti-Kv2.1 antibody in SHSY5Y cells expression Kv6.1WT but not the Met425 variant.

**(B)** SH-SY5Y cells were co-transfected with Kv6.1-HA (wild-type) and either Kv6.2-Flag (WT) or Kv6.3-Flag (WT). Kv6.1 was immunoprecipitated using an anti-HA antibody, and co-immunoprecipitated Kv6.2 or Kv6.3 was probed using an anti-Flag antibody. The co-immunoprecipitation results show that Kv6.1 does not interact with either Kv6.2 or Kv6.3. Notably, a weak interaction between Kv6.1 and Kv6.2 was observed in a single replicate (data not shown). *Input = lysate from transfected cells; IP = co-immunoprecipitated sample. N= number of replicates

