## Supplementary material for "A pain-reducing Kv6.4 variant spares other Kv6 channels, offering a target for uterine pain": Supplementary material1.pdf

**A**

| | | Primer Sequence (5' to 3') |
|---|---|---|
| **rs140124801** | KCNG1-F | GTCGCCATAGCCCATCGTCGTCATGGTGA |
| | KCNG1-R | TCACCATGACGACGATGGGCTATGGCGAC |
| | KCNG2-F | TCGCCGTAGCCCATGGTGGTCATGGAG |
| | KCNG2-R | CTCCATGACCACCATGGGCTACGGCGA |
| | KCNG3-F | CATATCTCCATAGCCCATTGTAGTCATAGAGATAATCACCCACCAG |
| | KCNG3-R | CTGGTGGGTGATTATCTCTATGACTACAATGGGCTATGGAGATATG |
| | KCNG4-F | GACATGGTGCCCCGCAGTGTGCCAGGC |
| | KCNG4-R | GCCTGGCACACTGCGGGGCACCATGTC |
| **rs140682724** | KCNG4-F | CCGTAGCCCACCATTGTCATGGAGATGATGGCC |
| | KCNG4-R | GGCCATCATCTCCATGACAATGGTGGGCTACGG |

**B**

| | Primer Sequence (5' to 3') |
|---|---|
| KCNG1-F | AGAACGAGATGGCCGACAG |
| KCNG1-R | CGGGAGAAGGTGTGGAAGAT |
| KCNG2-F | GACTTCTCCAGCGTGCCC |
| KCNG2-R | TGTGGAAGATGGAGGTGACC |
| KCNG3-F | TCAGCTTCTTGAACATGGGC |
| KCNG3-R | CTGACAACACAAACTCCTCCA |
| KCNG4-F | TCACCCTCTTCTCCCCTTTG |
| KCNG4-R | ATGATGAGGATCCCGCTCAG |
| KCNB1-F | GCTTTCGCTTTCTGCCTCG |
| KCNB1-R | GCTGTAGTCATCGCACACCT |

### Supplementary Fig S1

| ICD-10 CODE | Clinical phenotype |
|---|---|
| I10 | Essential (primary) hypertension |
| E78.0 | Pure hypercholesterolaemia |
| K57.3 | Diverticular disease of large intestine without perforation or abscess |
| K44.9 | Diaphragmatic hernia without obstruction or gangrene |
| J45.9 | Asthma |
| K21.9 | Gastro-oesophageal reflux disease without oesophagitis |
| Z87.1 | Personal history of diseases of the digestive system |
| Z86.7 | Personal history of diseases of the circulatory system |
| R07.4 | Chest pain |
| I25.1 | Atherosclerotic heart disease |
| H26.9 | Cataract |
| Z92.1 | Personal history of long-term (current) use of anticoagulants |
| M17.9 | Gonarthrosis |
| E66.9 | Obesity |
| E03.9 | Hypothyroidism |
| K59.0 | Constipation |
| F32.9 | Depressive episode |
| K63.5 | Polyp of colon |
| D64.9 | Anaemia |
| N39.0 | Urinary tract infection |
| M19.9 | Arthrosis |
| I25.9 | Chronic ischaemic heart disease |
| K29.7 | Gastritis |
| I20.9 | Angina pectoris |
| R10.4 | Other and unspecified abdominal pain |
| N40 | Hyperplasia of prostate |
| R11 | Nausea and vomiting |
| K40.9 | Unilateral or unspecified inguinal hernia |
| K44.9 | Diaphragmatic hernia without obstruction or gangrene |
| R07.4 | Chest pain |
| M17.9 | Gonarthrosis |
| Z96.6 | Presence of orthopaedic joint implants |
| M19.9 | Arthrosis |
| R10.4 | Other and unspecified abdominal pain |
| M16.9 | Coxarthrosis |
| R10.1 | Pain localised to upper abdomen |
| M13.9 | Arthritis |
| R51 | Headache |
| R10.3 | Pain localised to other parts of lower abdomen |
| M15.9 | Polyarthrosis |
| M54.5 | Low back pain |
| R07.2 | Precordial pain |
| M10.9 | Gout |
| M54.9 | Dorsalgia |
| M19.99 | Arthrosis (Site unspecified) |
| G43.9 | Migraine |
| M79.66 | Pain in limb (Lower leg) |
| M19.97 | Arthrosis (Ankle and foot) |
| M25.55 | Pain in joint (Pelvic region and thigh) |
| M06.9 | Rheumatoid arthritis |
| M19.91 | Arthrosis (Shoulder region) |
| M54.3 | Sciatica |
| G40.9 | Epilepsy |
| F03 | Unspecified dementia |
| G62.9 | Polyneuropathy |
| G30.9 | Alzheimer's disease |
| G20 | Parkinson's disease |
| F00.9 | Dementia in Alzheimer's disease |
| I69.4 | Sequelae of stroke |

**Supplementary Fig S2**

A

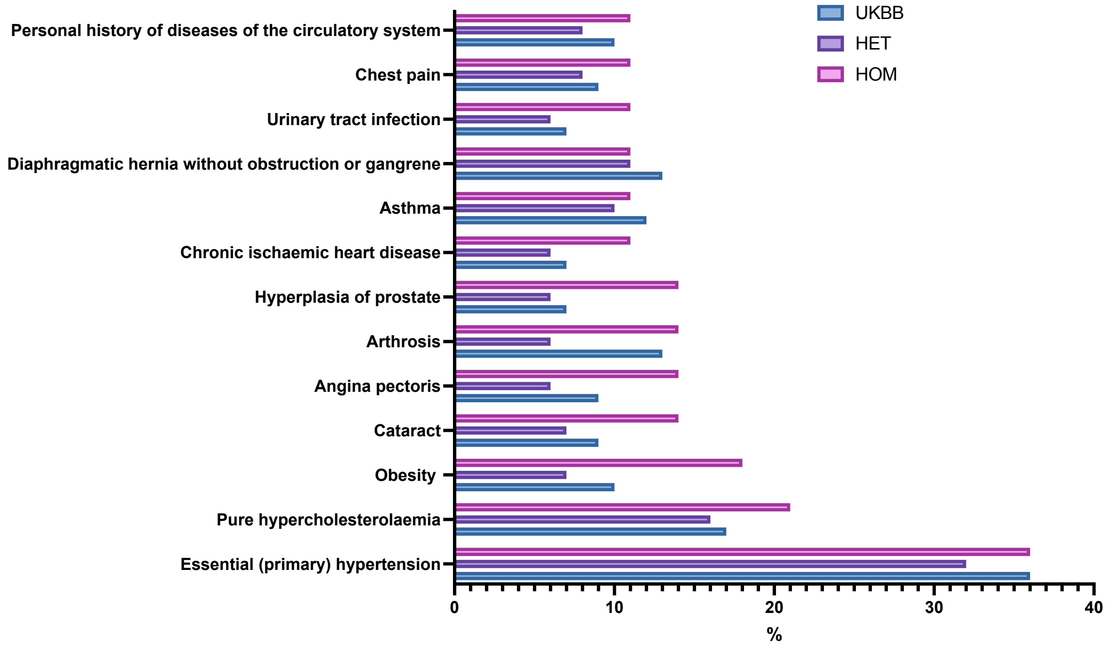

B

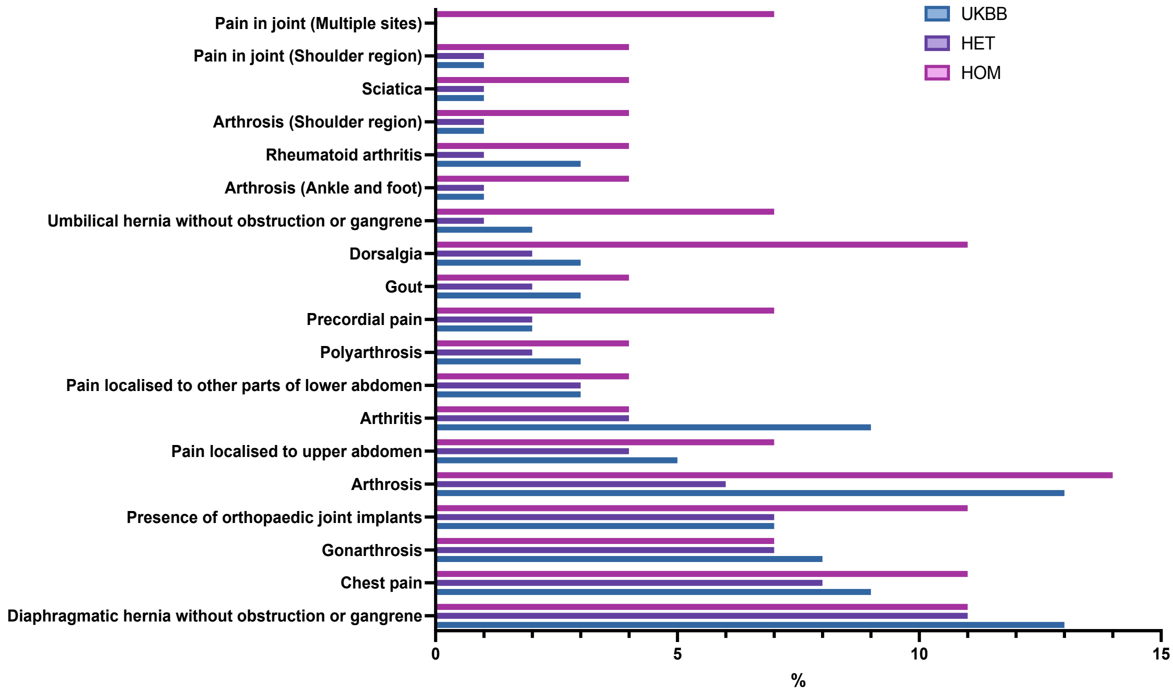

C

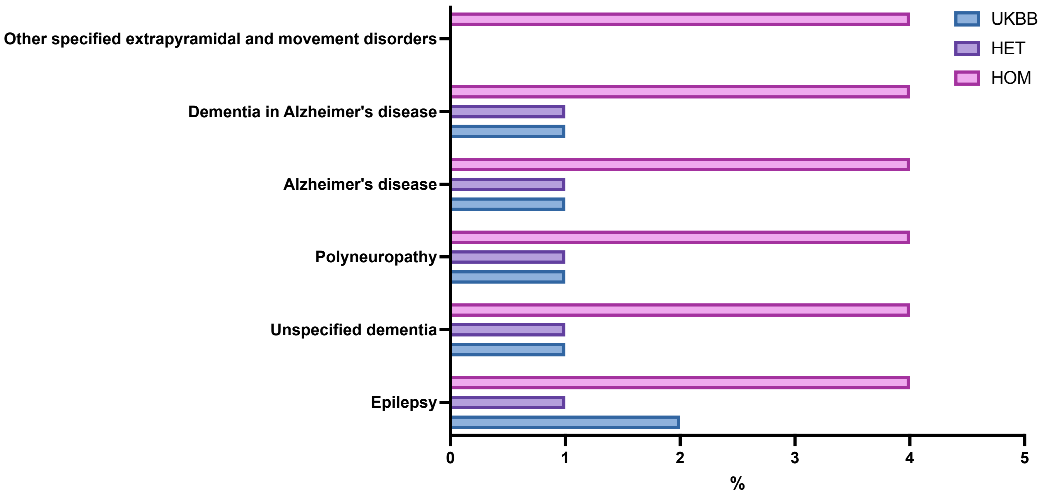

**Supplementary Fig S3**

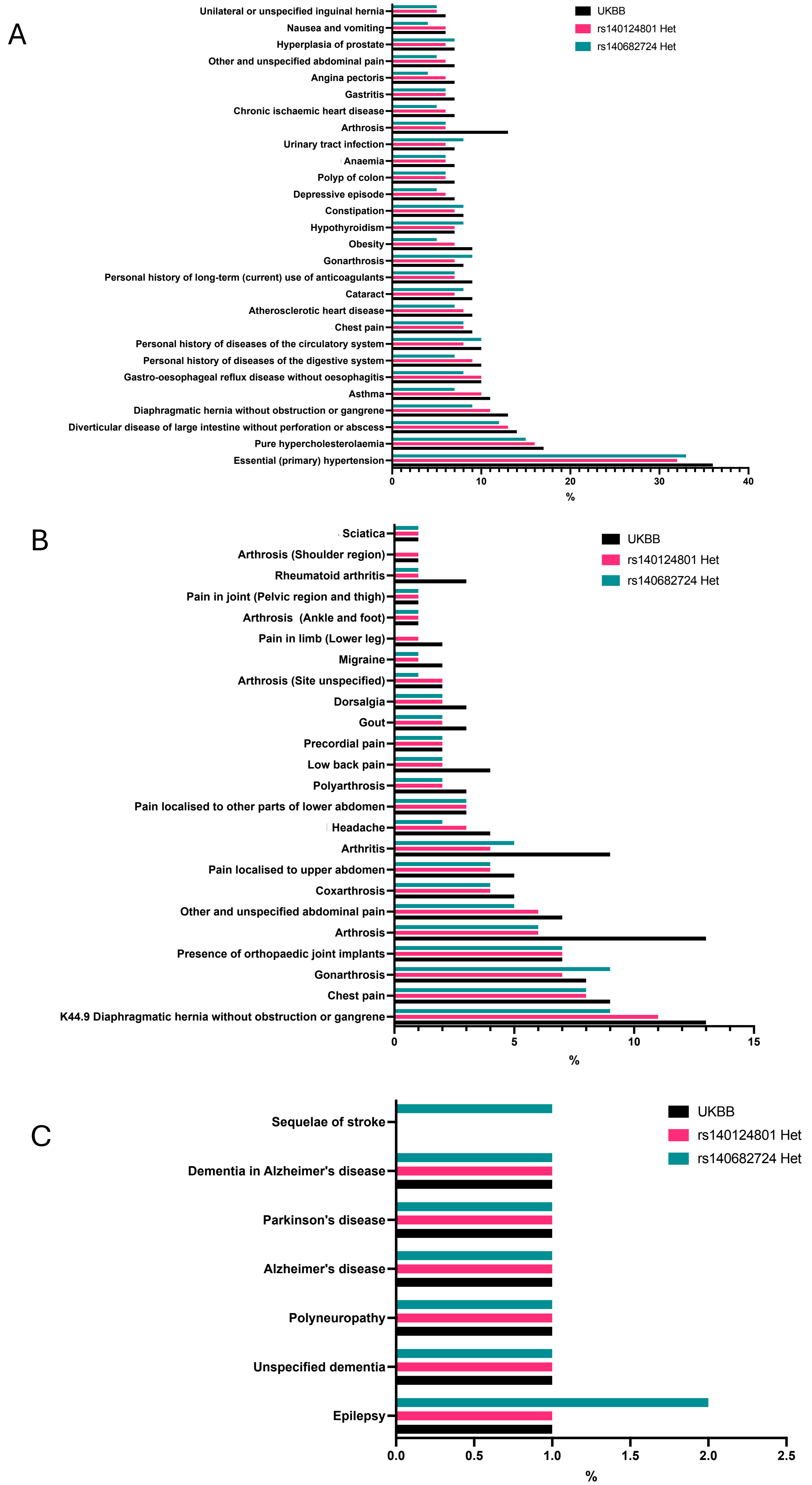

**Supplementary Fig S4**

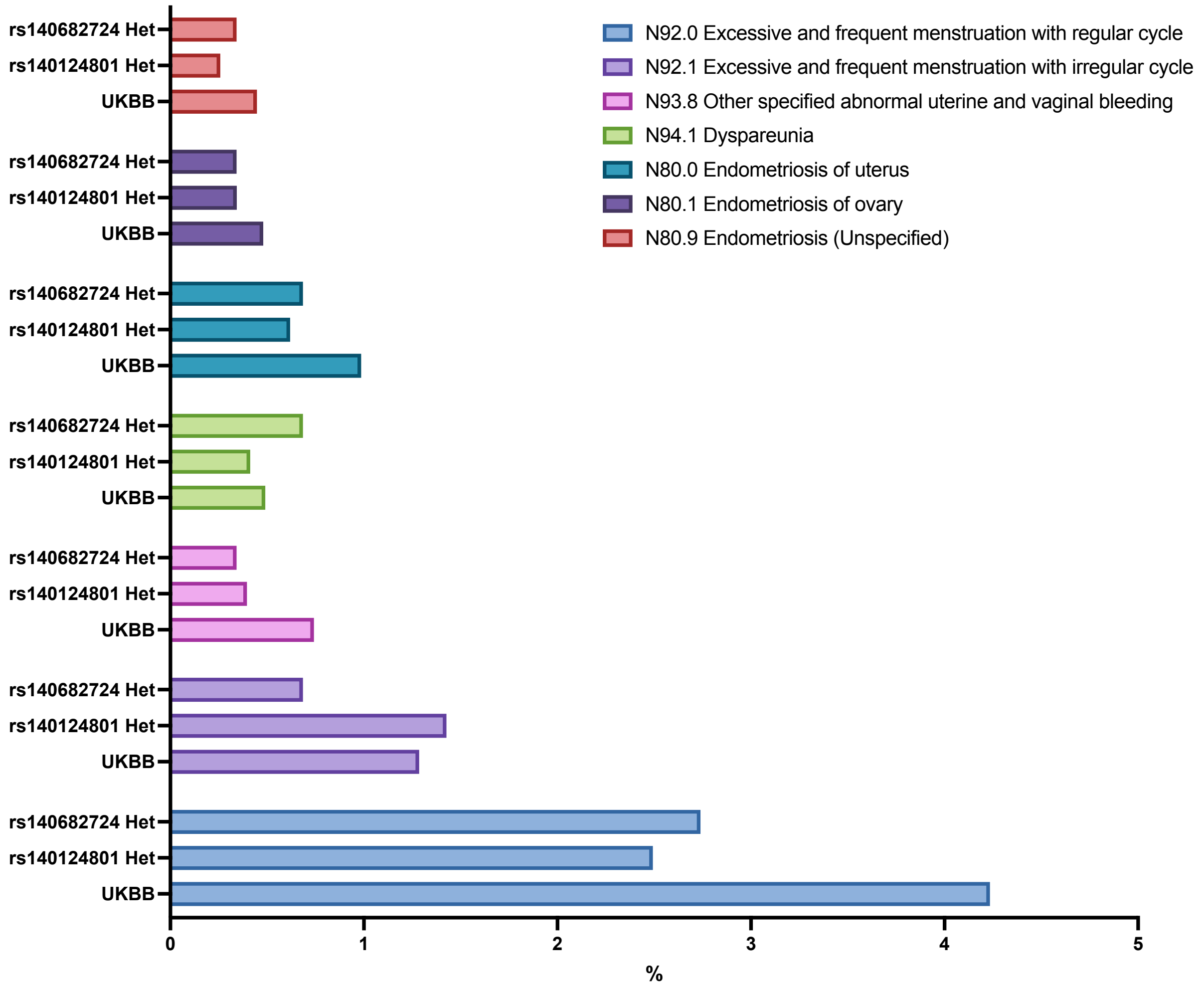

**Supplementary Fig S5**

**Supplementary methods:**

**qPCR:**

To quantify relative *KCNB1* mRNA levels in SHSY5Y and HEK293 cells, real-time PCR was performed by combining 5 ng of cDNA LightCycler® 480 SYBR Green I Master Mix (Roche Life Science, Penzberg, Germany), and 1 μM of both forward and reverse primers (primer sequences available upon request) in each reaction. Data were normalized to the housekeeping gene GAPDH. Samples were analyzed using a CFX384 Touch Real-Time PCR Detection System (Bio-Rad Laboratories). The associated software was used to calculate crossing threshold (Ct) values, and relative gene expression was subsequently calculated using the ΔΔCt method.

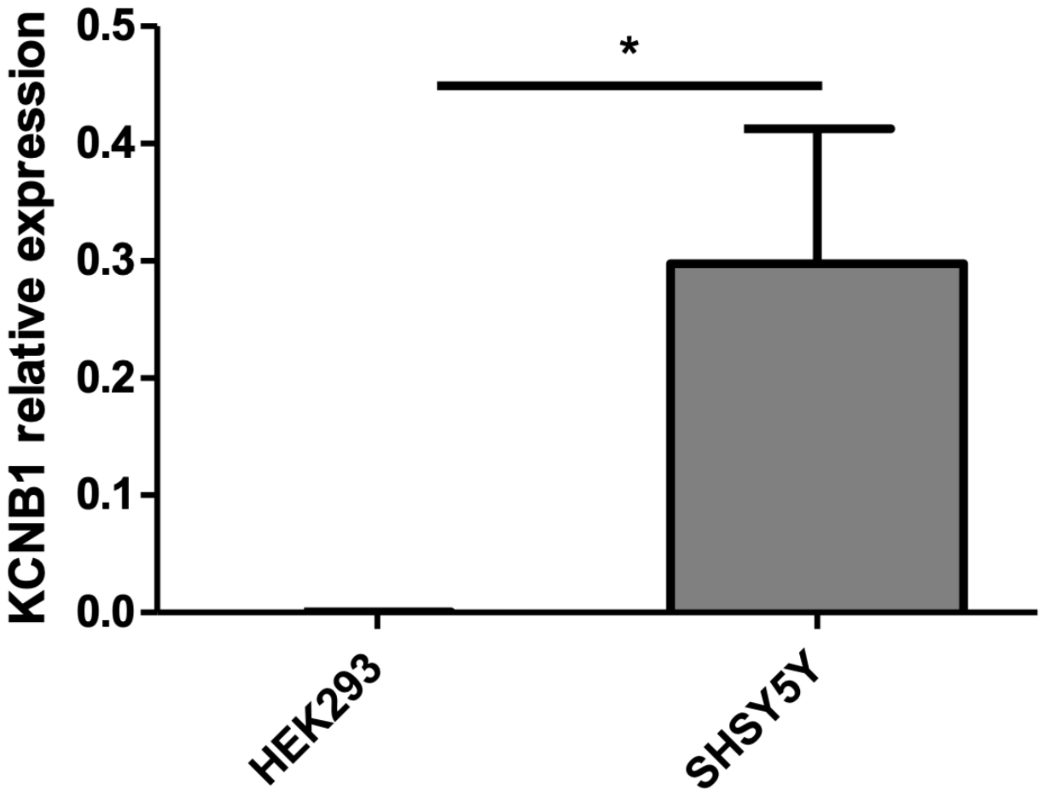

**Supplementary Fig S6**
